# Development of Biologically Interpretable Multimodal Deep Learning Model for Cancer Prognosis Prediction

**DOI:** 10.1101/2021.10.30.466610

**Authors:** Zarif L. Azher, Louis J. Vaickus, Lucas A. Salas, Brock C. Christensen, Joshua J. Levy

## Abstract

Robust cancer prognostication can enable more effective patient care and management, which may potentially improve health outcomes. Deep learning has proven to be a powerful tool to extract meaningful information from cancer patient data. In recent years it has displayed promise in quantifying prognostication by predicting patient risk. However, most current deep learning-based cancer prognosis prediction methods use only a single data source and miss out on learning from potentially rich relationships across modalities. Existing multimodal approaches are challenging to interpret in a biological or medical context, limiting real-world clinical integration as a trustworthy prognostic decision aid. Here, we developed a multimodal modeling approach that can integrate information from the central modalities of gene expression, DNA methylation, and histopathological imaging with clinical information for cancer prognosis prediction. Our multimodal modeling approach combines pathway and gene-based sparsely coded layers with patch-based graph convolutional networks to facilitate biological interpretation of the model results. We present a preliminary analysis that compares the potential applicability of combining all modalities to uni- or bi-modal approaches. Leveraging data from four cancer subtypes from the Cancer Genome Atlas, results demonstrate the encouraging performance of our multimodal approach (C-index=0.660 without clinical features; C-index=0.665 with clinical features) across four cancer subtypes versus unimodal approaches and existing state-of-the-art approaches. This work brings insight to the development of interpretable multimodal methods of applying AI to biomedical data and can potentially serve as a foundation for clinical implementations of such software. We plan to follow up this preliminary analysis with an in-depth exploration of factors to improve multimodal modeling approaches on an in-house dataset.

## 1 INTRODUCTION

### 1.1 Cancer and Cancer Prognosis

The global disease burden of cancer is far-reaching and is a major contributor to rising all-causes mortality and the global stagnation of life expectancy. In 2020 alone, an estimated 19.3 million new cancer cases were diagnosed worldwide, while nearly 10 million deaths were reported [29]. These staggering numbers underscore the imperative need for more effective therapies and treatment options. Consequently, cancer care has become increasingly precise and personalized, where medication and treatment for a patient are selected based on granular individualized data [17]. Researchers and practitioners are chiefly interested in ascertaining the disease prognosis from this information, which can indicate the risk of mortality over time by estimating a time-to-event outcome. Patient-specific risk scores provide critical insight for patient outcomes and care management and administration [31]. By nature, prognosis prediction is never certain, so, understandably, novel risk stratification methods from personalized data are constantly being researched.

### 1.2 Deep Learning for Cancer Prognosis Prediction

In recent years, artificial intelligence and machine learning technologies have shown promise for cancer prognosis prediction [1, 3, 14]. Traditional assessment methods quantify prognosis using Cox proportional hazards regression on a small set of clinical covariates [35]. These models assume proportional temporal dependence in the hazard (instantaneous risk) function between strata and that a linear dependence between a set of covariates and the log hazard function, as compared to baseline hazard, exists. Coefficients are estimated to represent the change in log hazard as a function of a one unit increase in the covariate. The hazard ratio numerically represents the instantaneous relative risk of an event occurring between strata. Most prognosis prediction models are configured to predict time to death. While traditional methods handpick clinical covariates such as age, weight, height, or other clinical factors, modern machine learning methods mine complex associations (e.g., interactions, nonlinear transformations) between high-dimensional input predictors/features and the outcome are generally impossible to curate manually. These approaches have been shown to be consistent with or outperform conventional approaches, spurring continued research of new methods [8, 15, 24, 32].

Machine learning methods for cancer prognosis prediction use a variety of data collected from patients. Omics data, including but not limited to gene expression or RNA sequencing, DNA methylation (DNAm), and copy-number variations (CNV), have become popular choices since they reflect early oncogenic effects indicative of metastatic potential. Recent multi-omics modeling approaches have sought to combine omics modalities as they are reflective of multiple levels of genetic and epigenetic co-regulation [33]. Genetic and epigenetic alterations may drive morphological and architectural presentations, observable with imaging modalities that capture tissue histology such as whole-slide imaging (WSI) from histopathological slides and radiological imaging (e.g., CT scans). For instance, histology slides are routinely used for disease staging (e.g., TNM staging system to assess local invasion, nodal and distant metastasis) [16], and this information may be combined with clinical information to communicate cancer survival. The resolution of these data types can vary widely. As an example, there are over 20,000 protein-coding genes with complex collinear/co-expression patterns, while WSI can store billions of elements but maybe reducible based on spatially redundant patterns.

Most approaches for prognosis prediction are unimodal because incorporating other data sources is either nontrivial or nearly impossible to acquire. However, recent multimodal modeling approaches have demonstrated improved predictive power [4, 6, 28, 30]. These multimodal models extract meaningful feature embeddings (descriptive n-dimensional vectors) of individual data sources and then combine these representations to output a hazard ratio prediction. Numerous neural network architectures have been developed to accomplish such tasks, typically operating on different input modalities. For example, Chen et al. [6] used a graph-convolutional network (GCN) and a convolutional neural network (CNN) for WSI, a standard feedforward artificial neural network (ANN) for gene expression, and an attention-based mechanism to fuse extracted embeddings. Another group [30] applied ANNs to clinical data, mRNA, miRNA, DNAm, and CNV, CNNs for representative patches of WSI, and a row-wise maximum method for embedding fusion. While there is no consensus on ideal multimodal deep learning architectures, generally, novel approaches will be bottlenecked by their capacity to model individual modalities while flexibly leveraging complementary information.

### 1.3 Interpretability of Deep Learning for Cancer Prognosis Prediction

Across both single-modality and multimodal methods for cancer prognosis prediction, most models suffer from a lack of interpretability – it is difficult to determine why the models make the predictions that they do. Furthermore, network architecture designs are largely selected based on experimental results alone and are not intuitively explainable to medical practitioners. For example, deep neural networks (DNNs), which are widely used in biomedical machine learning applications, can become a “black box,” which is difficult for practitioners to understand intuitively as layers are added [26]. The potential for real-world adoption suffers as a result, as a lack of understandability erodes trust and the likelihood of adoption for machine learning tools [27].

This phenomenon has prompted the development of interpretable neural network approaches for DNA methylation (DNAm) and gene expression [12]. While most neural networks for tabular data are fully connected, these networks utilize prune connections between layers to establish sparse/local connections between CpGs and genes or genes and pathways. Gene expression information is inputted, with data from each node/gene being transmitted to nodes in the next layer representing functional biological pathways related to specific genes. For imaging data such as WSI, attention-based approaches assign importance scores to regions of the slide corresponding to a positive prediction. GCNs have the added advantage of capturing nested spatial dependencies within tissue sections (complex macro-architectural relationships; e.g., tumor invading into deeper layers). To our knowledge, no multimodal deep learning approaches for cancer prognosis have been developed, which integrate both sparse local connections for omics information and graphs for histology.

### 1.4 Contributions

Here, we present a biologically interpretable multimodal deep learning framework for cancer prognosis prediction, which integrates gene expression and DNAm using locally connected neural networks, WSI with graph neural networks, and clinical covariates. Our work displays the potency of these interpretable network architecture choices and contributes to the understanding that multimodal data integration benefits cancer prognosis prediction. This study serves of a preliminary analysis of the potential utility of such software, which we plan to expand upon in future work.

## 2 METHODS

### 2.1 Data Acquisition

Gene expression, DNAm, WSI, and corresponding clinical covariates (e.g., age, race or ethnicity, sex, and TNM stage) for 1160 patients diagnosed with cancer were downloaded from The Cancer Genome Atlas (TCGA), representing primary tumors in colon (colorectal) (n=307), bladder (n=221), skin (melanoma) (n=435) and liver (n=405) (**Table 1**; **Table A.2.1**). Out of the 1160 total patients for which we had downloaded data, 952 were placed into the training and validation sets, while 208 were randomly placed into held-out testing sets using the train_test_split function available in the scikit-learn Python package using scikit-learn version 0.24.2 and Python version 3.7.10.

**Table 1:** Breakdown of training, validation and test samples

|  | Colon | Bladder | Skin | Liver | Totals |
| --- | --- | --- | --- | --- | --- |
| **Train** | 217 | 154 | 318 | 294 | 835 |
| **Val** | 39 | 28 | 57 | 53 | 117 |
| **Test** | 51 | 39 | 60 | 58 | 208 |
| **Total** | 307 | 221 | 435 | 405 | 1160 |

Clinical predictors were transformed into feature vectors. One-hot encoding was applied to generate a numerical representation of categorical features, including disease stage, patient race or ethnicity, sex, and primary cancer site. These one-hot encoded representations were input into the feature vectors without any further preprocessing. We did not represent stage as an ordinal variable (e.g., orthogonal polynomials).

### 2.2 Unimodal Gene Expression and DNAm Models

#### 2.2.1. Data Preprocessing for Gene Expression and DNAm

Methylation arrays were preprocessed using the workflow established through PyMethylProcess [23]. Using the supplementary DNAm annotation file, CpGs which are not associated with any genes were removed, as were non-autosomal and SNP-related CpGs (inclusion of such CpGs can introduce potential for bias which is beyond the scope of this work) [10, 34, 36], ultimately yielding 225,343 CpGs.

For gene expression and DNAm data, dictionary mappings were created and stored to map genes and CpGs to associated biological pathways and associated genes, respectively [21]. Gene set information was downloaded from MSigDB (C6 collection). Genes that were not associated with any MSigDb pathways were removed, leaving 10,868 genes.

#### 2.2.2. Sparse Pathway Layers for DNAm and Gene Expression and DNAm

For the gene expression ANN, each input node represents gene expression normalized by Fragments per Kilobase of transcript per million mapped reads (FPKM) (10,868 genes present across all patients in the datasets). This information is routed to corresponding signature oncogenic gene set signatures, which serve as nodes in the subsequent layer (pathway layer; 189 nodes).

For the DNAm ANN, each input node represents a CpG’s beta value, which is the proportion of methylated alleles at that site. Information from CpGs is routed to genes in the subsequent layer; CpG-gene mappings were configured using the Illumina 450K array annotations, downloaded from the Gene Expression Omnibus (GEO), analogous to previous investigations of sparse CpG-gene connections (19,025 genes) [21].

The input and a second biologically representative layer serve as encoder layers that extract meaningful gene expression and DNAm features. Additional standard fully connected layers are configured for ancillary and downstream (e.g., multimodal) tasks to improve feature extraction.

#### 2.2.3. Unsupervised Pretraining with Sparse Variational Autoencoders

We trained variational autoencoders (VAEs) using sparse connectivity (e.g., CpG to gene; gene to pathway) to initialize the encoder layers of our unimodal gene expression and DNAm models (**Figure 1**). Variational autoencoders learn to compress information with maximal fidelity to the original data source based on a reconstruction loss (mean-squared error; MSE; compares decoded information from latent distribution to original information), and the posterior distribution is penalized using the Kullback–Leibler (KL) divergence in order to conform to a multivariate normal distribution, which serves as a generative approach to regularize downstream training through the sampling of the posterior. VAEs are popular approaches for representing nuanced omics information and reducing data requirements of downstream supervised objectives if properly tuned [13]. The VAEs featured for the gene expression and DNAm unimodal models utilize sparse connections assigned via gene-pathway-gene and CpG-gene-CpG mappings, respectively, where the latent pathway and gene layers undergo reparameterization trick in order to differentiate through a posterior parameterized by a mean and variance (**Figure 1A**). Both VAE models used the Adam optimizer with a learning rate of 0.008, and were trained for 600 epochs.

**Figure 1:**
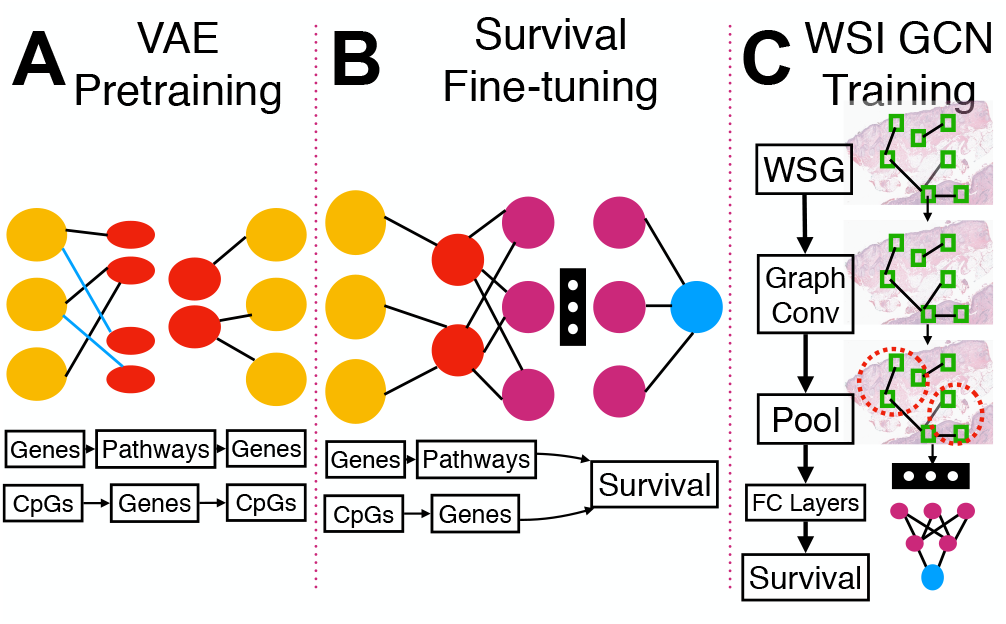
Graphical overview of unimodal modeling approaches: A) Gene expression and DNAm encoders are initially trained using sparsely coded VAEs; B) These encoders are further tuned during the training of the unimodal UniGE and UniDNAm models; C) The unimodal UniGCN is trained using survival targets.

#### 2.2.4. Fine-*tuning Unimodal Models with Survival Objectives*

Once the gene expression and DNAm VAEs were pre-trained, we reconfigured the encoder architecture to predict patient survival, transferring the previously learned encoder weights as initialization for the ANNs (**Figure 1B**). The encoder layers served as upstream feature extraction layers, from which hidden layers and an output layer were appended in order to output a hazard ratio prediction. Both of the models were trained using the Cox partial likelihood loss function.

The gene expression survival model (UniGE) was trained for 100 epochs using the Adam optimizer with a learning rate of 0.001 and a batch size of 24. The DNAm survival model (UniDNAm) was trained for 50 using the Adam optimizer with a learning rate of 0.001 and a batch size of 16.

### 2.3 Unimodal WSI Model

We created a graph convolutional network (GCN) – UniGCN – to predict cancer prognosis from WSI (**Figure 1C**). GCNs are particularly suited for WSI because convolutions across nodes of a graph, in contrast to Euclidean data, are invariant to permutation and can naturally capture complex macro architectural phenomena in tissue sections operating solely on subarrays of a WSI which exhibit spatial dependence [20]. To our knowledge, this work is one of the first applications of GCNs to WSI to predict cancer survival.

#### 2.3.1. Data Preprocessing for WSI

We configured the UniGCN to make predictions across spatially adjacent tissue patches in contrast to treating cells as nodes due to computational complexity constraints and the greater weight assigned to tissue macroarchitecture versus potentially missed information when relying solely on cells. WSI were split into 256×256 pixel patches to serve as nodes of the graph (denoted as a Whole Slide Graph, WSG), connected using a radius neighbors algorithm (pytorch_cluster) to define spatial adjacency. To form nodal attributes for each image patch, 2048-dimensional embedding vectors were extracted from the penultimate layer of a Resnet-50 CNN (pre-trained on the ImageNet dataset) using the PathFlowAI software package [19].

#### 2.3.2. Survival-Objective Graph Convolutional Network for Whole-Slide-Imaging Pretraining

We constructed a GCN to predict hazard ratios from WSG using the Pytorch Geometric library [9]. Information was passed amongst stochastically selected neighborhoods of nodes using SAGEConv convolutional operators while undergoing transformation to 100-dimensional vectors [11]. Two successive such layers were applied to further contextualize node embeddings by aggregating from neighboring nodes. Graph convolutions were followed by node normalization and ReLU activation functions. After the application of three such graph convolutional layers, information was pooled across nodes using the SAGPool operator, which assigns an attention score to each node to decide its importance for the downstream layer [18], where a fraction of nodes with low attention is pruned. After multiple applications of graph convolution and pooling layers, information from the remaining nodes were aggregated into a single vector of length 100 to represent the entire WSI using a GlobalAttention layer. Finally, fully connected layers were applied to predict a hazard ratio.

UniGCN was trained for 20 epochs, and the parameters were updated using a learning rate of 0.008 with the Adam optimizer. The training batch size was 10.

### 2.4 Multimodal Model for Prognosis Prediction

Embeddings extracted from each of the penultimate encoder layers of the unimodal survival models (UniGE, UniDNAm, UniGCN). The multimodal prognosis model aggregates these unimodal embeddings via either concatenation or averaging to form an aggregate omics-WSI embedding which encapsulates all supplied input modalities (**Figure 2**). In addition, a 2-layer multilayer perceptron (MLP) encodes clinical covariate predictors and adds this information to the aggregate multimodal embedding information. The aggregate embedding information is then transformed using fully connected layers and an output layer to predict a hazard ratio. In contrast to other multimodal modeling frameworks, this framework is initialized with two identical pre-trained unimodal encoders for each modality (**Figure 2A-B**), trained as specified in sections 2.2-2.3. One set of encoders – the *Similar Arm* – is intended to capture information shared across the three central modalities – gene expression, DNA methylation, and WSI – while the other one – Different Arm – aims to create embeddings encoding features distinct to each modality (**Figure 2B)**. In each arm, encoders from unimodal survival models and the clinical encoder extract embeddings. Then, these embeddings are projected to a common dimension or length.

**Figure 2:**
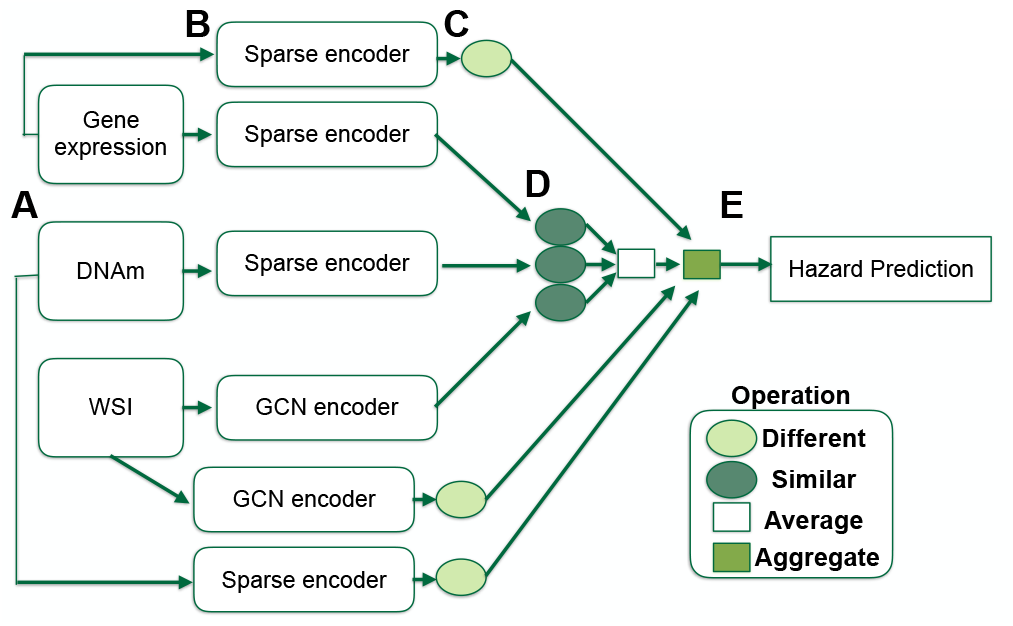
The double-arm multimodal prognostication model. Information from A) each modality is passed through B) modality-specific encoders, producing embeddings which are C) Different and D) Similar. Similar embeddings are averaged, while Different embeddings are E) combined with similar information to predict hazard. Representations are aggregated and used to make a prognosis prediction. Not pictured is the clinical encoder, which is present in each arm in ClinMulti and StagedMulti.

The model averages embeddings from the encoders meant to generate similar embeddings (*Similar Arm*; **Figure 2C**), while it concatenates embeddings from the encoders intended to create different embeddings (*Different Arm*; **Figure 2D**). Next, the concatenated *Different Arm* representations is then merged with the averaged *Similar Arm* representation and passes through two fully connected output layers to generate a hazard ratio for each patient or sample (**Figure 2E**).

The multimodal model is trained using the Cox partial likelihood loss, with the Adam optimizer and a learning rate of 0.0008. The learning rate was derived using experimentation across values of 0.001, 0.0008, and 0.0001. We trained three variations of the multimodal model; *StagedMulti*, which used disease stage as a clinical variable and other clinical features; *ClinMulti*, which excluded disease stage from the clinical features; and *NormMulti*, which did not use the clinical modality at all. We created these delineations because clinical information such as staging can provide valuable prognosis-related information, while we wish to primarily assess the power of our model to extract information from molecular and WSI modalities. *NormMulti* was trained for 16 epochs with a batch size of 5, *ClinMulti* was trained for 14 epochs with a batch size of 3, and *StagedMulti* was trained for 12 epochs with a batch size of 3. Gradient accumulation was used for *ClinMulti* and *NormMulti* to accommodate for the relatively small batch sizes.

### 2.5 Model Evaluation and Comparison

Compared to our multimodal cancer prognosis model, we trained and tested different survival analysis approaches on the same datasets. For both the clinical model including stage as a covariate and excluding stage, we trained a gradient-boosted Cox proportional hazards model (GradCoxPH), a random forest model (RF), and an ElasticNet-penalty Cox proportional hazards model (ElastiCox) [25]. Two versions of each model were trained; one which excluded disease staging information (prefix Lim-), and one where disease staging information was included (prefix Full-). These models had patient sex as an additional variable. The multimodal clinical models did not utilize sex due to observed overfitting. All of comparison models were implemented using the scikit-survival Python package.

Additionally, we compared the results of the multimodal approaches to the aforementioned unimodal DNAm, gene expression, and WSI survival models, as well as a prognostic model trained on solely omics modalities – DNAm and gene expression – and a final model trained on all central modalities, which used a single encoder arm, concatenated representations, and used fully connected layers, instead of the *Similar* and *Different Arms* implemented for our central multimodal models (SingleMulti). These ensured that we could validate our multi-arm scheme and justify the use of our modalities and model architectures.

We updated the model parameters using the training set for all modeling approaches and employed early stopping procedures/coarse hyperparameter scanning utilizing the validation set. Across each cancer subtype, we evaluated the performance of our prognosis prediction models on the test set using the Concordance-Index (C-index), a common metric used to evaluate survival models that compares the assigned risk score and event timepoints via a rank correlation. C-index values range from 0.0 - 1.0, where a score of 0.5 indicates random predictions, while 1.0 indicates perfect survival predictions. We report C-indices specific to the following cancer subtypes – colon (Colon Adenocarcinoma), bladder, skin (Melanoma), and liver. C-indices were averaged across subtypes to estimate a macro-averaged (across subtype) C-index, and C-indices were also estimated across the test cohort, irrespective of subtype, to yield an Overall C-index. We estimated 95% confidence intervals using 1000-sample nonparametric bootstrapping to assess statistical significance for the model comparisons.

### 2.6 Interpretation Methods

#### 2.6.1. Interrogating important genes and pathways for omics models with permutation feature importance

To interrogate which genes or pathways were important to the UniDNAm and UniGE, we permuted nodes from the gene and pathway layers 100 times. Permutation of gene or pathway values is expected to reduce the C-statistic. We report the average reduction in C-index across permutations of the gene/pathway as its feature importance [2]. Such importances were ranked to display gene/pathways, which were necessary for the model prediction.

#### 2.6.2. Important WSG nodes with Self-Attention Graph Pooling

The WSI GCN encoder features self-attention layers which assign importance scores to patches for inclusion in a subsequent layer. Thus, patches which remain after the application of each pooling layer are successively more important. We extracted and visualized image patches that were deemed to be most important by the model by recording those that remained after applying each graph pooling layer. We chose to use the WSI GCN encoder from the NormMulti *Similar Arm* for this analysis.

## 3 RESULTS AND DISCUSSION

### 3.1 Survival Results

#### 3.1.1. Unimodal Survival Results

The gene expression network achieved an overall C-index of 0.653, while both the DNAm neural network reached a C-index 0.616, and the WSI GCN reached an overall C-index of 0.616 (**Table 2**).

**Table 2:**
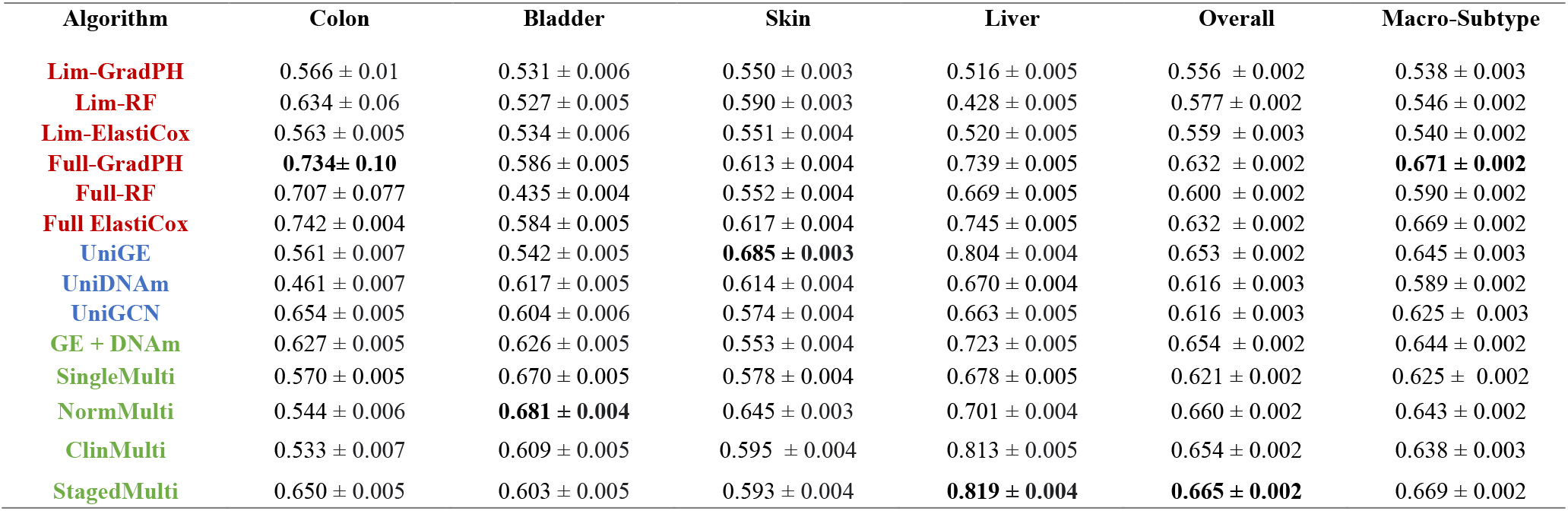
Survival Analysis C-Indices: Algorithms are colored by whether they use: A) Clinical covariates alone (red), B) Single Omics/WSI modality (Blue), c) More than one modality (Green)

| Algorithm | Colon | Bladder | Skin | Liver | Overall | Macro-Subtype |
| --- | --- | --- | --- | --- | --- | --- |
| **Lim-GradPH** | 0.566 ± 0.01 | 0.531 ± 0.006 | 0.550 ± 0.003 | 0.516 ± 0.005 | 0.556 ± 0.002 | 0.538 ± 0.003 |
| **Lim-RF** | 0.634 ± 0.06 | 0.527 ± 0.005 | 0.590 ± 0.003 | 0.428 ± 0.005 | 0.577 ± 0.002 | 0.546 ± 0.002 |
| **Lim-ElastiCox** | 0.563 ± 0.005 | 0.534 ± 0.006 | 0.551 ± 0.004 | 0.520 ± 0.005 | 0.559 ± 0.003 | 0.540 ± 0.002 |
| **Full-GradPH** | **0.734 ± 0.10** | 0.586 ± 0.005 | 0.613 ± 0.004 | 0.739 ± 0.005 | 0.632 ± 0.002 | **0.671 ± 0.002** |
| **Full-RF** | 0.707 ± 0.077 | 0.435 ± 0.004 | 0.552 ± 0.004 | 0.669 ± 0.005 | 0.600 ± 0.002 | 0.590 ± 0.002 |
| **Full ElastiCox** | 0.742 ± 0.004 | 0.584 ± 0.005 | 0.617 ± 0.004 | 0.745 ± 0.005 | 0.632 ± 0.002 | 0.669 ± 0.002 |
| **UniGE** | 0.561 ± 0.007 | 0.542 ± 0.005 | **0.685 ± 0.003** | 0.804 ± 0.004 | 0.653 ± 0.002 | 0.645 ± 0.003 |
| **UniDNAm** | 0.461 ± 0.007 | 0.617 ± 0.005 | 0.614 ± 0.004 | 0.670 ± 0.004 | 0.616 ± 0.003 | 0.589 ± 0.002 |
| **UniGCN** | 0.654 ± 0.005 | 0.604 ± 0.006 | 0.574 ± 0.004 | 0.663 ± 0.005 | 0.616 ± 0.003 | 0.625 ± 0.003 |
| **GE + DNAm** | 0.627 ± 0.005 | 0.626 ± 0.005 | 0.553 ± 0.004 | 0.723 ± 0.005 | 0.654 ± 0.002 | 0.644 ± 0.002 |
| **SingleMulti** | 0.570 ± 0.005 | 0.670 ± 0.005 | 0.578 ± 0.004 | 0.678 ± 0.005 | 0.621 ± 0.002 | 0.625 ± 0.002 |
| **NormMulti** | 0.544 ± 0.006 | **0.681 ± 0.004** | 0.645 ± 0.003 | 0.701 ± 0.004 | 0.660 ± 0.002 | 0.643 ± 0.002 |
| **ClinMulti** | 0.533 ± 0.007 | 0.609 ± 0.005 | 0.595 ± 0.004 | 0.813 ± 0.005 | 0.654 ± 0.002 | 0.638 ± 0.003 |
| **StagedMulti** | 0.650 ± 0.005 | 0.603 ± 0.005 | 0.593 ± 0.004 | **0.819 ± 0.004** | **0.665 ± 0.002** | 0.669 ± 0.002 |

Overall C-indices for unimodal comparison approaches (GradPH, RF, ElasticCox) are reported in **Table 2**.

#### 3.1.2. Multimodal Survival Results

The GE+DNAm approach achieved an overall C-index of 0.654. SingleMulti, which is identical to NormMulti except it uses a single encoder arm, scored an overall C-index of 0.621. NormMulti, which did not use any clinical features, achieved an overall C-index of 0.660, while ClinMulti, which additionally used clinical features of age, race, and cancer subtype, achieved an overall C-index of 0.654. Finally, StagedMulti, which used age, race, cancer subtype, and disease stage, scored an overall C-index of 0.665 (**Table 2**). ClinMulti likely achieved a lower C-index than NormMulti despite having access to a greater quantity of data, due to an inability to make sense of the extra data without the added stage information.

### 3.2 Model Interpretation

#### 3.2.1. Identified pathways from gene expression model

In table **A.3.1** we list the 5 most impactful gene sets (ranked from highest to lowest) as determined by permutation importance by subtype and overall for the gene expression model. The pathways with the highest permutation importance are potentially prognostically relevant biomarkers as determined by our model.

#### 3.2.2. Identified genes from DNAm model

In table **A.3.2** we list the 5 most impactful genes (ranked from highest to lowest) as determined by permutation importance by subtype and overall for the DNAm model. DNA methylation-associated genes which our model identified may be relevant for cancer prognostication (**A.3.2**).

#### 3.2.3. Salient tissue regions identified by multimodal GCN model

Visual inspection of salient tissue regions as determined by the GCN pooling layers of the Similar Arm in the NormMulti model, pointed to potential interactions between tumor and tumor-infiltrating immune cells, in one example as related to better survival (**Figure 3A**). This evidence further supports that the GCN model is picking up potentially prognostically relevant regions rather than random noise (**Figure 3**).

**Figure 3:**
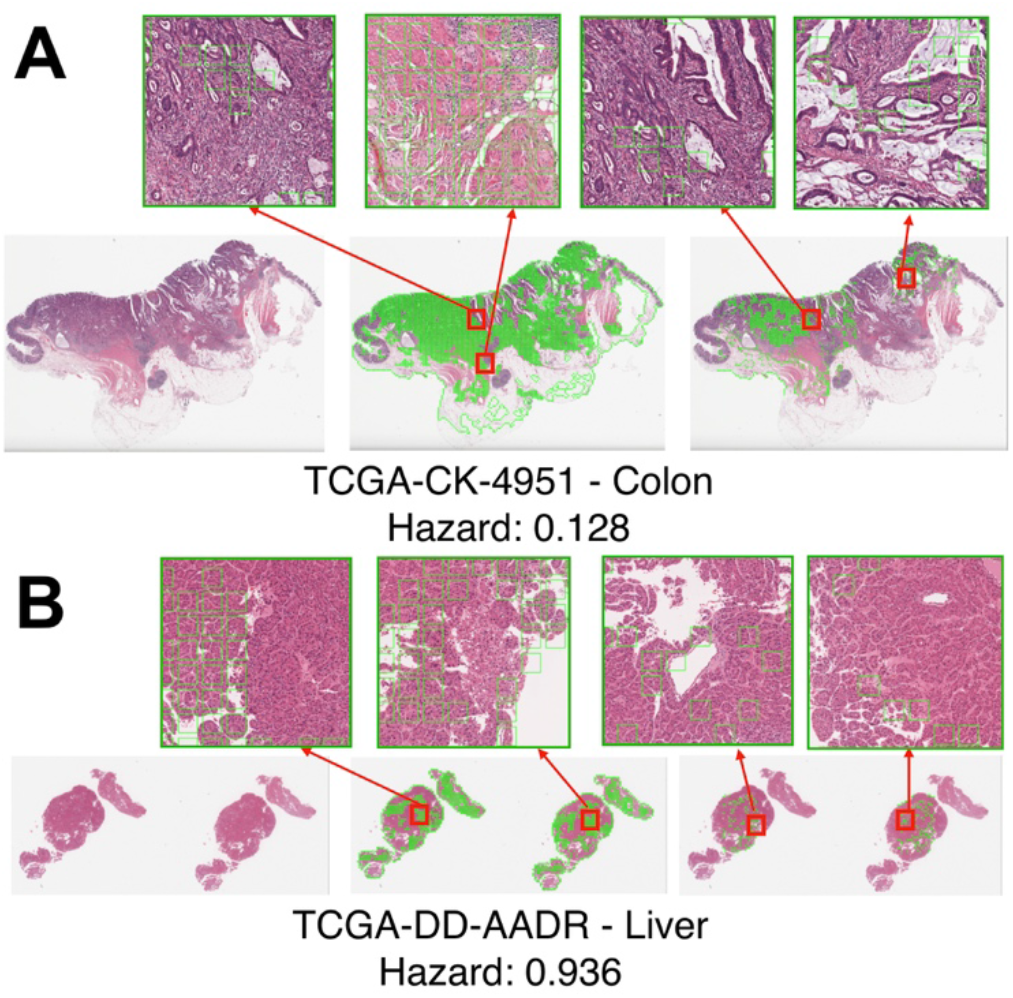
Visualized patches given to WSI segments by pooling layer. As the WSI graph is progressively pooled, differing regions of the image selected by the model are given attention and are thus deemed potentially relevant for prognostication: A) TCGA-CK-4951 was predicted to be low risk by the GCN, while B) TCGA-DD-AADR was predicted to be high risk.

### 3.3 Discussion

This study sought to develop multimodal modeling approaches that could leverage complementary information from distinct clinical, omics, and imaging modalities to improve prognostic, predictive capacity versus modeling a single modality by itself. Results from our study demonstrate that the overall model performance for multimodal approaches is greater than using a single modality at a time. Furthermore, we found that adding clinical information, particularly the TNM stage, improves predictive performance.

Our method compares favorably to other multimodal approaches, which also purport to have built-in interpretability. For instance, our NormMulti model outperforms the state-of-the-art PORPOISE [5] bladder, liver, and skin (8.6%, 6.1%, and 9.5% on bladder, skin, and liver cancer, respectively) while featuring analogous interpretation mechanisms. Other multimodal survival models (e.g., MultiSurv) achieve remarkable performance but have limited interpretation since they do not feature GCNs, attention, or local connections, which can elucidate such findings [30]. Given these results, we believe that we have potentially established a relevant modeling framework for multimodal cancer prognosis, especially for bladder, skin, and liver cancer.

The DNAm sparse survival model and the WSI GCN performed similarly among unimodal models when considering the entire testing dataset, including all cancer subtypes. Notably, our WSI GCN, with a bladder-specific C-index of 0.616, statistically significantly outperformed Patch-GCN (C-index of 0.585) [7] for bladder survival prediction - the only other existing WSI GCN for cancer prognostication, to our knowledge. The gene expression model achieved the highest C-index out of any unimodal model. Overall, utilizing clinical predictors alone, without staging information, was suboptimal to DNAm, gene expression, and WSI. However, adding TNM clinical stage of colon cancer tumors to the set of clinical predictors, as utilized by the gradient boosted Cox proportional hazards model, achieved the highest colon-specific C-index of all trained models for colon, which is unsurprising since staging criteria in the colon is well defined and captures well the tradeoffs between local invasion and nodal/distant metastasis (METS). Using the GCN alone on WSI, we would expect to largely capture signals pertaining to local invasion only, with some indication of the potential for METS (e.g., immune infiltration; **Figure 3A**). Since results from WSI GCN and the staging-based model were similar in the colon, this is suggestive that inspection of the biopsy at the initial invasion site can help understand the route of metastasis, though nuanced molecular information may provide further confirmation [22]. However, clinical methods and models which incorporated stage (e.g., StagedMulti performed worse than NormMulti and ClinMulti) performed significantly poorer than other models for cancer types other than colon, suggesting a greater reliance on complementary molecular information. We plan to investigate these findings further to better understand the histological correspondence of local/distant METS and the optimal alignment of clinical staging scales for prognostication based on this information.

### 3.4 Limitations

Although this work features a preliminary analysis of multimodal prognostic models, we plan to fine-tune our approach to address potential study limitations and improve the generalizability of these approaches for application in external cohorts. For instance, the relatively small and somewhat class-imbalanced dataset we curated may have contributed to poor model performances in some subtypes such as bladder cancer. Coarse optimization of hyperparameters, including learning rate, batch size, hidden layer sizes, etc., may have also contributed to suboptimal performance. Both of these constraints may be ameliorated through additional data collection and the adoption of modeling approaches that can reduce study bias (e.g., Bayesian methods, which can enforce a prior over the neural network weights to reduce the potential for spurious findings). Additional penalizations supplied to the model parameters present an opportunity to explore targeted hypotheses (e.g., concordance between modalities, direct identification of similar and disjoint components of variation).

Another real-world element that our model does not cover is cases of missing data when a patient who needs prognostication does not have data available for all modalities. This can be addressed using techniques such as training with missing data scenarios through the adoption of attention mechanisms or adversarial robustness measures, as demonstrated in previous works [4]. Lower-resourced areas may not have access to quality omics and/or imaging data in the global health context, where these mechanisms may prove handy.

Finally, our gene expression network is limited by the MSigDB C6 biological pathway collection, as only genes present in this set are included in our network. Investigating different organizations of capsules may allow our network to learn critical prognostication features and test the prognostic relevance of select gene sets or pathways.

### 3.5 Future Directions

Though our multimodal approaches yielded the highest overall performance, there is much room for improvement for specific subtypes. Given that different unimodal models may excel in different subtypes, we plan to explore the role of attention mechanisms in dynamically up weighting more relevant omics/imaging modalities and why such information was up weighted for a particular patient. Furthermore, different methods for pretraining (e.g., predicting expression from DNAm to initialize ANN weights) may improve unimodal encoders, which may improve the robustness of multimodal modeling methods, as initialized parameters for multimodal approaches may be more informative. For example, instead of using a ResNet-50 CNN model pre-trained on ImageNet to extract WSI patch features, contrastive predictive coding (e.g., self-supervised methods for WSI patches that capture surrounding architecture) may improve results. Furthermore, self-supervised methods for graphs exist but have not been well explored in the context of histology. Further work will investigate the impact of less commonly used modalities such as radiological imaging, additional objective functions applied to models, transfer learning of models between different cancer subtypes, and orthogonal validation of salient biomarkers.

We plan to conduct a follow-up internal validation study at the Dartmouth-Hitchcock Medical Center, which will envision the potential applicability of multimodal software while implementing additional method improvements.

## 4 CONCLUSIONS

We presented a biologically interpretable multimodal deep learning model framework to predict cancer patient prognosis, which is the first to integrate DNAm, gene expression, and WSI in a way that may optimally realize the dependencies amongst predictors (e.g., genes to pathways, WSI as graphs). We demonstrated the capacity of these multimodal modeling to extract potential disease-relevant pathways while reporting in important tissue subcomponents with graph pooling. Based on comparisons with existing methods, our multimodal approach has the potential to be established as a state-of-the-art method in interpretable cancer prognosis prediction for certain cancer subtypes, pending additional fine-tuning of the approach. This study sheds light on the positive potential of explainable neural network design for cancer prognosis prediction and the power of integrating multiple modalities into decision making.

## APPENDIX

### A.2 Methods

#### A.2.1 Dataset Covariate Statistics

|  |  | Subtype |  |  |  |
| --- | --- | --- | --- | --- | --- |
|  |  | Colon | Bladder | Skin | Liver |
| **Number of Samples** |  | 307 | 221 | 435 | 405 |
| **Race, number (%)** | Asian | 10 (3.1) | 29 (12.6) | 12 (2.5) | 168 (39.2) |
|  | Black or African American | 65 (20.1) | 13 (5.6) | 1 (0.2) | 24 (5.6) |
|  | Not reported | N/A | 12 (5.2) | N/A | N/A |
|  | White | 232 (76.9) | 167 (76.6) | 422 (97.3) | 213 (55.2) |
| **Sex, number (%)** | F | 148 (47.8) | 55 (23.8) | 163 (38.8) | 144 (33.6) |
|  | M | 159 (52.2) | 166 (76.2) | 272 (61.2) | 261 (66.4) |
| **Stage, number (%)** | Stage 0 | N/A | N/A | 7 (1.5) | N/A |
|  | Not Reported | 8 (2.5) | 2 (0.9) | N/A | 34 (7.9) |
|  | Stage i | 48 (14.8) | N/A | 77 (16.2) | 173 (45.9) |
|  | Stage ii | 113 (39.5) | 73 (31.6) | 136 (29.5) | 96 (22.4) |
|  | Stage iii | 89 (27.5) | 76 (35.1) | 155 (36.5) | 96 (22.4) |
|  | Stage iv | 49 (15.1) | 70 (32.5) | 24 (5.1) | 6 (1.4) |
| **Age, mean (SD)** |  | 50.7 (16.3) | 68.2 (10.8) | 50.3 (20.2) | 60.8 (20.0) |

### A.3 Results and Discussion

#### A.3.1 Identified Potential Pathway Biomarkers; Most Important Pathways - Highest to Lowest

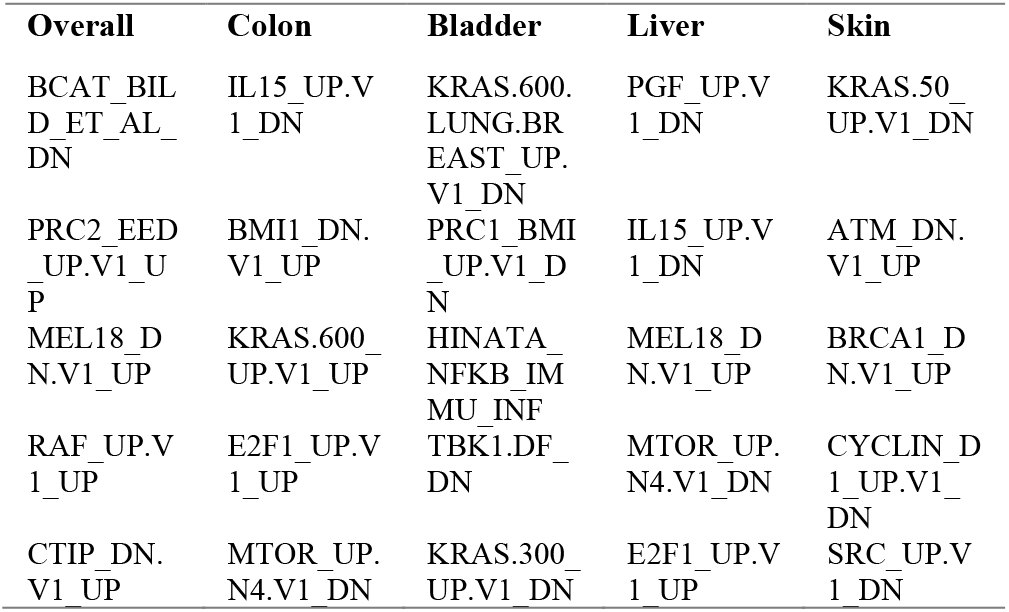

#### A.3.2 Identified Potential Gene Biomarkers; Most Important Genes - Highest to Lowest

| Overall | Colon | Bladder | Liver | Skin |
| --- | --- | --- | --- | --- |
| LZTS2 | CIB1 | TMED2 | PCSK2 | TRIOBP |
| ZCWPW1 | MCOLN2 | ZNF71 | FUT6 | BMT2 |
| OR10H1 | LINC00596 | DRD5 | HSD11B2 | PLA2G4B |
| CHI3L1 | MIF | MIR1258 | CAPNS1 | TBPL1 |

| CKLF | SSH1 | MILR1 | SLC5A11 | ACIN1 |
| --- | --- | --- | --- | --- |

